# PAR recognition by PARP1 regulates DNA-dependent activities and independently stimulates catalytic activity of PARP1

**DOI:** 10.1101/2021.12.21.473685

**Authors:** Waghela Deeksha, Suman Abhishek, Eerappa Rajakumara

## Abstract

Poly(ADP-ribosyl)ation is predominantly catalyzed by Poly(ADP-ribose) polymerase 1 (PARP1) in response to DNA damage, mediating the DNA repair process to maintain genomic integrity. Single strand (SSB) and double strand (DSB) DNA breaks are bonafide stimulators of PARP1 activity. However, PAR mediated PARP1 regulation remains unexplored. Here, we report ZnF3, BRCT and WGR, hitherto uncharacterized, as PAR reader domains of PARP1. Surprisingly, these domains recognize PARylated protein with a higher affinity compared to PAR but bind with weak or no affinity to DNA breaks as standalone domains. Conversely, ZnF1 and ZnF2 of PARP1 recognize DNA breaks but weakly to PAR. In addition, PAR reader domains, together, exhibit a synergy to recognize PAR or PARylated protein. Further competition binding studies suggest that PAR binding releases DNA from PARP1, and WGR domain facilitates the DNA release. Unexpectedly, PAR showed catalytic stimulation of PARP1 but hampers the DNA-dependent stimulation. Altogether, our work discovers dedicated high-affinity PAR reader domains of PARP1 and uncovers a novel mechanism of allosteric stimulation, but retardation of DNA-dependent activities of PARP1 by its catalytic product PAR. Therefore, our studies can be used as a model to understand the effect of one or more allosteric activators on the regulation of receptors or modular enzyme activities by another allosteric activator.

---

Modular proteins are highly regulated by diverse orthosteric (binds to active site) and/or allosteric (binds to other than active site) ligands to tune their functioning in the cell^1–6^. Binding of allosteric modulators to these enzymes exerts non-uniform effects on their functioning^7–14^. Many a times, the catalytic product(s) of the enzymes themselves act as an important regulator^5,10,12,15–17^. Therefore, exploring the allosteric regulation of enzymes would help not only to understand how signaling cross-talk fine-tunes physiology or cellular functioning but also to develop safer and more selective therapeutics^6–9^. In the current work, we explore the allosteric regulation of a modular enzyme, ADP-ribosyltransferase Diptheria toxin-like 1 (ARTD1) also known as Poly(ADP-ribose) polymerase 1 (PARP1), by its catalytic product Poly-ADPribose (PAR), and effect of PAR on DNA-dependent allosteric regulation of PARP1 catalytic activity.

PARP1 is an abundant nuclear enzyme often referred to as a “genome guardian” which plays a crucial role in maintaining genome integrity^18^. It is involved in several cellular functions, including DNA damage repair^19^, regulation of chromatin structure^20^, transcription^20,21^, and apoptosis^22,23^. PARP1 is a multi-domain protein consisting of three zinc-finger (ZnF) domains, ZnF1, ZnF2, and ZnF3 at the N-terminal, followed by the BRCT (Breast cancer associated C-terminal) domain and WGR (Trp-Gly-Arg rich) domains in the central region, and catalytic (CAT) domain at the C-terminal, which comprises a helical sub-domain (HD) and a signature ADP-ribosyl transferase (ART) sub-domain (Figure 1a). PARP1 has been shown to recognize different types of damaged DNA by different combinatorial sets of ZnFs. The ZnF1, ZnF3 and WGR together bind to double strand break (DSB), whereas ZnF1 and ZnF2 bind to single strand break (SSB) DNA and induces transient interaction of ZnF3 with DNA^24,25^.

Further, recognition of damaged DNA by DNA binding domains of PARP1 induces a global conformational change in the protein, i.e., spatial re-organization of the domains, which facilitates the positioning of CAT domain close to the automodification (AD) region, which comprises the C-terminal region of BRCT domain and the loop connecting the BRCT and WGR domains^24,26–28^. PARP1, then, catalyzes a series of covalent addition of ADP-ribose (ADPr) on the AD region using NAD^+^ as a co-factor to form a long, linear or branched, chain of up to >200 units of negatively charged ADP-ribose (ADPr), often referred to as PAR, and the process being known as PARylation^24,29^ (Figure 1b). PARP1 catalyzes the PARylation activity not only on itself (self-PARylation) but also on the other target proteins (trans-PARylation) and the target residues include aspartate, glutamate, lysine, serine, arginine, etc.^30–32^.

The enzymatic activity of PARP1 is allosterically regulated by different DNAs^24,33–35^. Additionally, PARP1 is also known to be activated upon binding to different types of RNA like snoRNA, pre-mRNA and therefore involved in RNA biogenesis^36,37^, alternative splicing of mRNA^38^, etc. However, the degree of activation of PARP1 upon RNA binding is less compared to DNA binding^39^. Proteins such as histones^40^, HPF1^1^, mH2A1.1, HMGN1, XPA, NEIL1, OGG1, DDB2, p53, ERK2, Sam68, YB-1, C12orf48, etc^2^ physically interact with PARP1 and regulate its catalytic activity. Histone PARylation factor (HPF1) protein, which forms a joint active site with PARP1, acts as a regulatory factor to promote serine PARylation over other target residues^1,30,41,42^. In addition, the interaction of PARP1 with other proteins or another PARP1 molecule is strengthened significantly in presence of DNA or PAR owing to the involvement of additional protein domains, for example, recognition of PAR on PARP1 by C-terminal domain (CTD) of p53 allows PARP1 to recognize and PARylate p53 at specific site^43^. Among several regulatory factors, the role of PAR in PARP1 regulation has not been explored extensively.

PAR serves as a docking platform for recruitment of DNA damage repair proteins. Several domains in various proteins have been identified to bind PAR. Besides the classical, well characterized PAR reader modules, macro domain, WWE, PBZ (PAR binding Zinc finger), PBM (PAR binding motif), from PAR hydrolyzing enzymes, there are other domains which have been reported to recognize PAR. These include FHA (Fork head associated) domain, OB (oligonucleotide or oligosaccharide binding fold), BRCT, etc.^44,45^.

Given that the chemical nature of PAR polymer is similar to nucleic acid molecules, like RNA/DNA (Figure S1), and PARP2, another DNA regulated enzyme of the PARP family, was reported to recognize and regulated by PAR^39^, we raised the following questions: (a) how does PARP1 recognize PAR? Whether the DNA break recognizing domains or other domains of PARP1 are involved in PAR recognition? (b) does PAR compete with DNA for PARP1 binding? (c) in case PARP1 recognizes PAR, does PAR regulate the PARP1’s catalytic activity? and (d) how does PAR binding affect the DNA dependent catalytic activity of PARP1?

Here, using biophysical approaches, we established the role of different domains of PARP1, individual and in combination, in recognition of PAR and PARylated protein (parProtein). Our studies show that although both PAR and DNA (SSB and DSB) are nucleic acid biopolymers, PARP1 recognizes them distinctly: high-affinity PAR binding domains bind weakly to DNA and vice versa. Our work also reveals that PAR acts as an allosteric stimulator of PARP1 activity and also partially inhibits its DNA dependent catalytic activity. Further, we demonstrate that PAR binding facilitates DNA dislodging from PARP1.

## Results and Discussion

### Biophysical characterization of PAR binding to PARP1

Recently reported EMSA study has shown that PARP1 binds to PAR^46^. Here, we determined the kinetics of PAR binding to PARP1 using biolayer interferometry (BLI). For the BLI study, we used biotinylated PAR and purified recombinant PARP1 (Figure S2a). Our study showed that PAR binds to PARP1 with an equilibrium molar dissociation constant (K_D_, a.k.a. binding affinity) of ∼39 nM (Figure 1c). The association rate constant (k_on_) and the dissociation rate constant (k_off_) for PAR binding to PARP1 were ∼5×10^4^ M^-1^ s^-1^ and ∼2×10^−3^ s^-1^, respectively (Figure 1c). The nanomolar of K_D_ and slower k_off_ suggest that PAR binds tightly to PARP1 and forms a stable complex. We further investigated the domains of PARP1 involved in the recognition of PAR.

**Figure 1.**
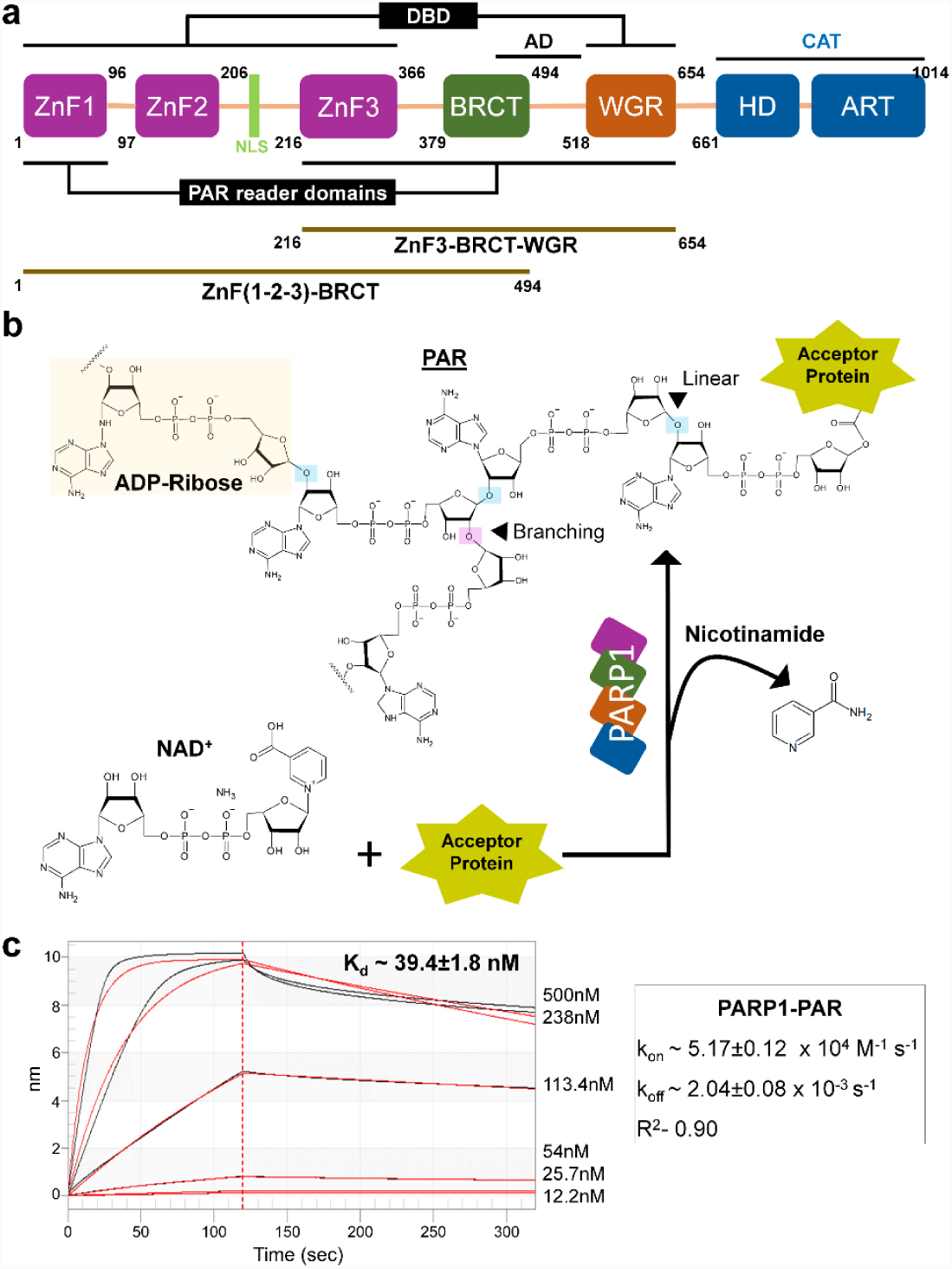
Domain architecture of PARP1 and PAR binding studies. **a** Schematic representation of the domain architecture of human PARP1 with dark brown bars below the architecture represents the multi-domain constructs of PARP1 used in the study. Black bars above and below the representation indicate their binding preferences. DBD, AD and CAT stands for ‘DNA binding domain’, ‘auto-modification domain’ and ‘catalytic domain’, respectively. **b** Representation of PARylation reaction catalyzed by PARP1 with chemical structures of NAD^+^, nicotinamide, ADP-ribose and Poly(ADP-ribose). Black triangles indicate the site of linear polymerization and branching in PAR. **c** BLI sensogram for PARP1 binding to PAR. Data was fit globally to the 1:1 binding model. The processed experimental data curves are in black; model fit curves from the 1:1 global analysis are shown in red. The values on the right side of each data show the range of analyte (PARP1) concentrations used for the experiment. The binding kinetics data are given in the inset on the right.

### Binding studies of DNA-break binding domains of PARP1 to PAR

Zn-finger domains along with WGR domain of PARP1 have been reported to recognize DNA-break^24^. However, no data is available whether these domains recognize PAR. Here, we have performed ITC experiments to investigate the possibility of PAR recognition by the individual DNA-break binding domains of PARP1. We have used recombinant DNA-break binding domains (Figure S2b-e) and purified PAR (Figure S3) to characterize their binding. The ZnF1 domain of PARP1 (ZnF1_PARP1_) bound to PAR with a K_D_ of ∼150 μM (Figure 2a) while ZnF2 domain of PARP1 (ZnF2_PARP1_) bound weakly to PAR (K_D_ could not be determined) (Figure 2b). Compared to ZnF1, the binding affinity of ZnF3 of PARP1 (ZnF3_PARP1_) for PAR increased by a factor of ∼53 (K_D_ ∼ 3 μM) (Figure 2c).

**Figure 2.**
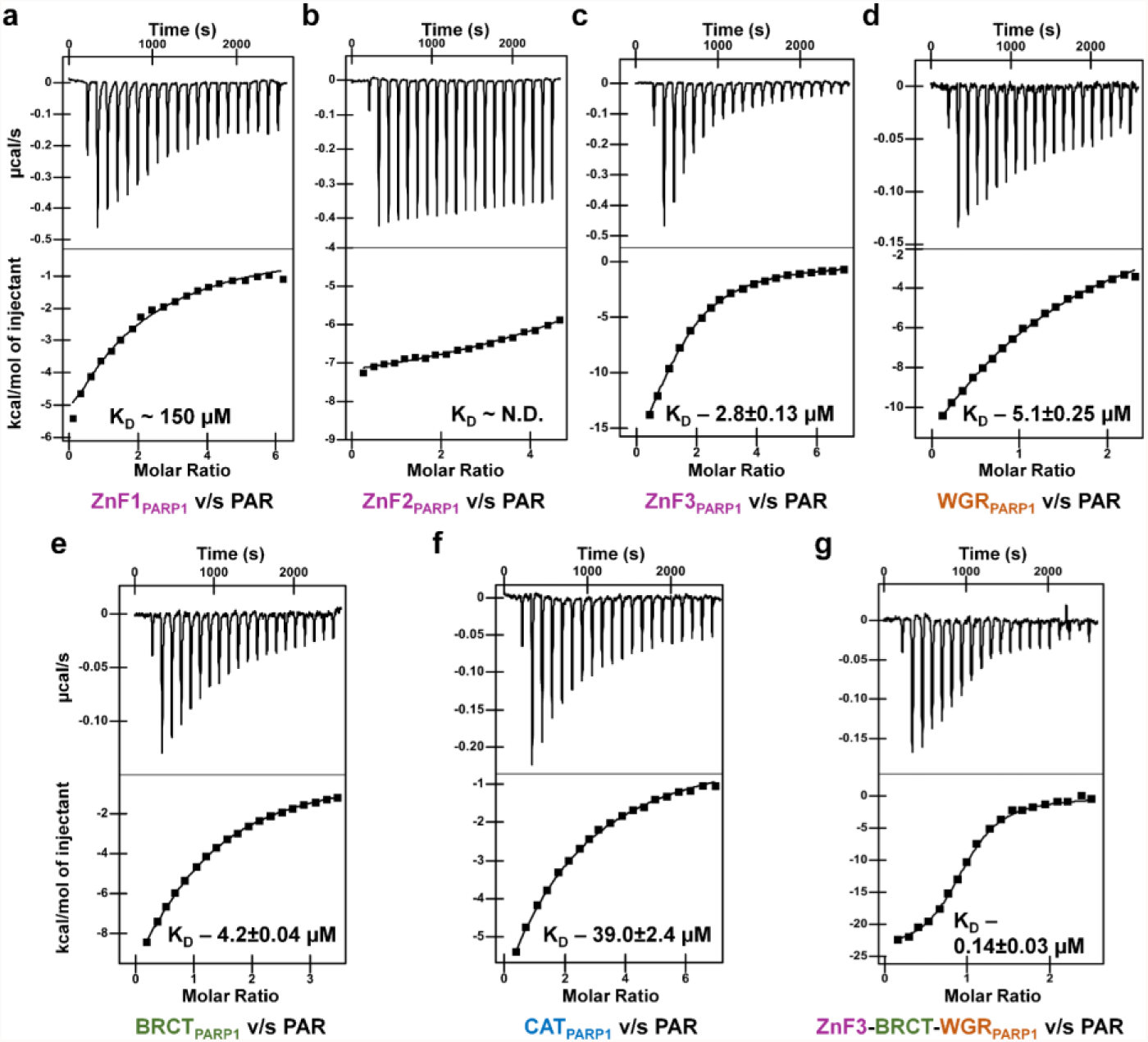
ITC measurements of PAR with domains of PARP1. Raw ITC data (upper panel) and normalized integration data (lower panel) for binding of PAR to **a** ZnF1_PARP1_, **b** ZnF2_PARP1_, **c** ZnF3_PARP1_, **d** WGR_PARP1_, **e** BRCT_PARP1_, **f** CAT_PARP1_, and **g** ZnF3-BRCT-WGR_PARP1_. K_D_ was determined using NanoAnalyze software and reported as mean ± s.e.m. (n=3).

Additionally, we recapitulated the previous findings that ZnF1_PARP1_ and ZnF2_PARP1_ have high affinity for DNA breaks (SSB and DSB DNA)^25,47,48^ (Figure S4a-d). Our further study show that ZnF3_PARP1_ exhibits weak binding to these DNAs (Figure S4e-f). These data show that Zn-fingers of PARP1 have different preferences for their binding partners. Structural comparison of Zn-fingers shows that ZnF3 is significantly different from ZnF1 and ZnF2 as it could not be superimposed on either of the two ZnFs, whereas ZnF1 and ZnF2 share high structural similarity with a RMSD of 0.6 Å for Cα atoms (Figure S5).

WGR domain of PARP1 (WGR_PARP1_), in combination with ZnF domains, has been known to recognize DSB DNA^24,28^, however WGR_PARP1_, as a standalone domain, doesn’t bind to DNA-break^49^. Here, we investigated the ability of standalone WGR_PARP1_ to recognize PAR. Surprisingly, the WGR_PARP1_ didn’t only bind to PAR but showed a high binding affinity with a K_D_ of ∼5 μM (Figure 2d). Together these binding studies demonstrate that among DNA-break binding domains of PARP1, ZnF3 and WGR bind to PAR with the highest affinity followed by ZnF1, whereas ZnF2 loosely binds to PAR. Also, as a control, we carried out buffer-PAR titration, which showed negligible heat of dilution (Figure S6). This study opens a window for the characterization of the WGR domain of PARP2 and PARP3 for the recognition of PAR, and its possible implications in DNA repair and epigenetic processes.

### BRCT and CAT domains of PARP1 also bind to PAR

The BRCT domains of proteins involved in DNA repair, such as XRCC1 (X-ray repair cross-complimenting protein 1) and ligase 4, have been shown to recognize PAR^45^. Considering it, we hypothesized that BRCT domain of PARP1 (BRCT_PARP1_) may also recognize PAR. We performed ITC binding studies of purified recombinant BRCT_PARP1_ (Figure S2f) with purified PAR (Figure S3). Our study showed that like ZnF3 and WGR domains, the BRCT_PARP1_ also exhibit high binding affinity (K_D_ ∼4 μM) for PAR (Figure 2e). Cryogenic electron microscopy structure (PDB ID: 7SCY, 7SCZ) reported that BRCT_PARP1_ bind to intact DNA in the nucleosomal context but no report on BRCT_PARP1_ binding SSB and DSB as standalone domain^50^. Taken together these binding studies suggest that three contiguous domains of PARP1, i.e., ZnF3, BRCT and WGR independently recognize PAR with high affinity. The current study supports the previous studies that the contiguous N-terminal domains of PARP1, i.e., ZnF1 and ZnF2, are high-affinity binders of DNA breaks. Here, for the first time, we have shown that PARP1 recognizes PAR with high affinity through its contiguous ZnF3, BRCT and WGR domains located at its central region (Figure 1a).

Since, the CAT domain of PARP1 (CAT_PARP1_) carries out the catalysis of PAR formation, therefore it might require recognizing the previous added unit(s) of ADPr for chain elongation. To explore this possibility, we used ITC to assess the binding of recombinant CAT_PARP1_ to purified PAR (Figure S2g and S3). Our study showed that CAT_PARP1_ exhibits a binding affinity of ∼39 μM for PAR (Figure 2f), which is ∼20 times weaker than that of ZnF3 domain. This study suggests that apart from amino acid substrate (target residue for ADP-ribosylation)^30–32^, CAT domain also recognizes the PAR subunits for further chain extension.

### Combinatorial recognition of PAR by PARP1 domains

As mentioned above, different combination of DNA-break binding domains of PARP1 cooperatively recognize SSB and DSB DNAs^24,25,48^. Here, we aim to assess the recognition of PAR by the identified high-affinity binder domains, ZnF3_PARP1_, BRCT_PARP1_ and WGR_PARP1_, in a single polypeptide over individual domains. To address this, we performed ITC binding study using recombinant ZnF3-BRCT-WGR_PARP1_ (Figure 1a, Figure S2h and S3), which showed that ZnF3-BRCT-WGR_PARP1_ binds to PAR with a K_D_ of ∼140 nM (Figure 2g). The binding affinity of ZnF3-BRCT-WGR_PARP1_ for PAR is ∼20, ∼30 and ∼36 times more than individual ZnF3_PARP1_, BRCT_PARP1_ and WGR_PARP1_ domains, respectively (Figure 2c-e & g), which suggest synergy among these domains to recognize PAR.

### Studies of PARP1 domains to recognize PARylated proteins

In cells, most PAR is present as a post-translational modification (PTM) on proteins, catalyzed by the PARP enzymes^51^, whereas, naked PAR is transiently present in the cell before it is completely degraded by PAR hydrolyzing enzymes^52^. Therefore, to explore the biological relevance, we hypothesize that the PAR reader domains of PARP1 may also recognize PAR on PARylated proteins. To test the hypothesis, we carried out ITC binding studies using PARylated PARP2 (parPARP2) with the identified PAR reader domains of PARP1, as standalone and in combination. For these studies, we performed *in vitro* PARylation of purified recombinant PARP2 (Figure S2i) to generate the parPARP2. Our studies showed that the ZnF3_PARP1_, BRCT_PARP1_ and WGR_PARP1_ bind to parPARP2 with a marginally higher affinity of ∼2, ∼2 and ∼3 μM, respectively (Figure 3a-c), compared to these domains binding to naked PAR (Fig. 2c-e). Further, titration of parPARP2 against ZnF3-BRCT-WGR_PARP1_ showed a K_D_ of 40 nM (Figure 3d), which is 3 times the affinity of construct binding to PAR (Figure 2g). These studies demonstrate that PARylated proteins are recognized with higher binding affinities by individual and combination of PAR reader domains, compared to naked PAR. To rule out the possibility that binding of these domains to PARP2 may contribute to enhancement in binding affinity, we carried out binding studies between these domains with PARP2 which showed that these domains do not bind to un-PARylated PARP2 (Figure S7). The higher binding affinity for PAR and PARylated protein by ZnF3-BRCT-WGR_PARP1_ compared to individual domains reconfirms that these domains show synergism for PAR recognition (Figure 2c-e, g and 3).

**Figure 3.**
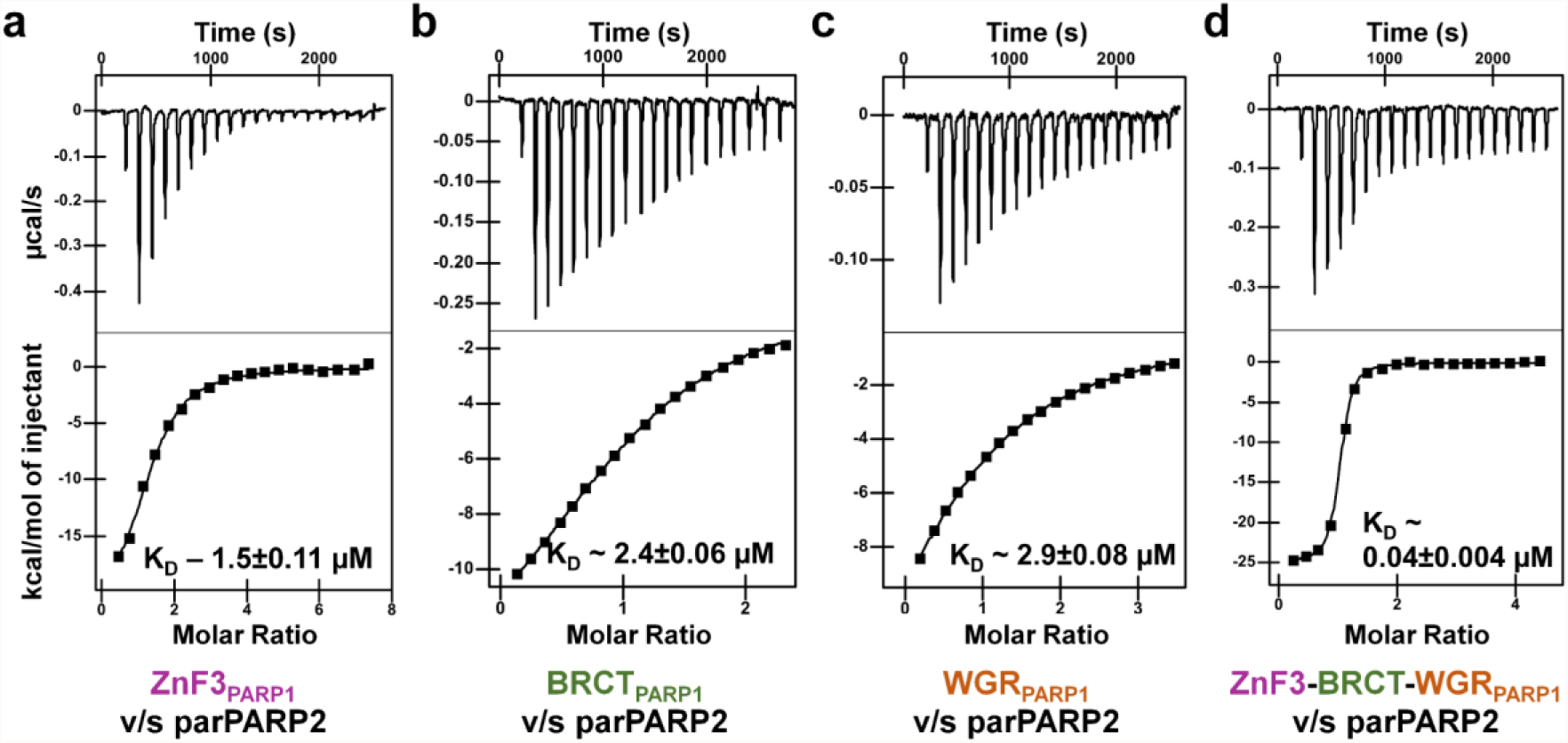
ITC measurements of PARylated PARP2 with domains of PARP1. Raw ITC data (upper panel) and normalized integration data (lower panel) for binding of PAR to **a** ZnF3 _PARP1_, **b** BRCT_PARP1_, **c** WGR_PARP1_, and **d** ZnF3-BRCT-WGR_PARP1_. K_D_ was determined using NanoAnalyze software and reported as mean ± s.e.m. (n=3).

### Studies on PAR induced dissociation of different DNA breaks from PARP1

It has been reported that upon self-PARylation, of all the DNA binding PARPs, i.e., PARP1, PARP2 and PARP3, release the bound DNA^33^. However, how the cross recognition of PAR, i.e., recognition of PAR on other PARylated proteins, by PARP1 affects the DNA-PARP1 binding dynamics is not known. To investigate the same, we carried out fluorescence polarization (FP) based competition binding studies of PAR and different DNA-breaks for PARP1 (Table S1, Figure S2a and S3). First, we performed FP binding studies of PARP1 with different DNA-breaks. PARP1 showed a K_D_ of ∼15 and ∼64 nM with DSB and SSB, respectively (Figure S8) as previously reported^25,35,48^. For competition FP assays, displacement of the fluorophore labelled bound DNA probe from PARP1 by PAR was evaluated. The molar equilibrium dissociation constant (*K*_*i*_) of the competitor PAR was determined to be ∼105 and ∼28, respectively, for DSB and SSB DNA bound to PARP1 (Figure 4). These competition binding studies delineate that PAR can effectively compete for both SSB and DBS breaks for binding to PARP1.

**Figure 4.**
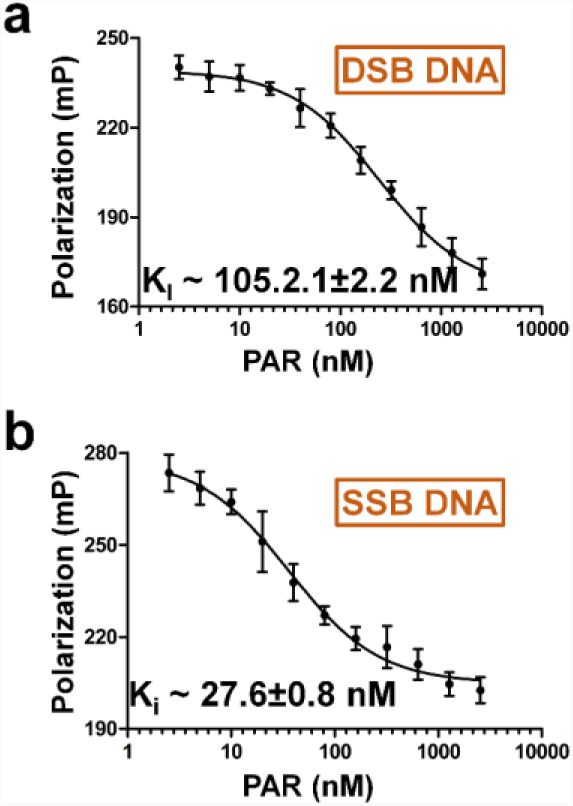
Fluorescence polarization-based competition binding study. Competition binding study of unlabeled PAR for binding to PARP1 against fluorophore labelled **a** DSB DNA and **b** SSB DNA. In FP experiments, PARP1 was incubated with either DSB or SSB before adding unlabeled PAR in increasing concentration. FP readings were measured after 30 minutes of incubation of the complexes. A decrease in polarization represents the replacement of 5-FAM labeled DNA by PAR. Error bar indicates standard deviation. All the data shown are from n=3 independent experiments. K_i_ is reported as mean ± s.e.m.

### Role of WGR domain in PAR induced DNA-break dissociation from PARP1

The WGR domain of PARP1, together with Zn-Finger domains, has been known to recognize damaged DNA^24,28^. Simultaneously, the above studies have identified WGR as a PAR reader domain (Figures 2d and 3c) and that PAR induces DNA dissociation from PARP1. This prompted to investigate the role of WGR domain in PAR-dependent dissociation of DNA break from PARP1. For that, we first determined the binding affinities of different DNA breaks for the recombinant variant of PARP1, ZnF1-ZnF2-ZnF3-BRCT (abbreviated as ZnF(1-2-3)-BRCT_PARP1_) (Figure S2j), which lacks WGR and CAT domains (Figure 1a). Our FP binding studies showed that ZnF(1-2-3)-BRCT_PARP1_ binds to DSB and SSB DNAs with a K_D_ of ∼88 and ∼70 nM, respectively (Figure 5a-b). Binding affinity of DSB DNA for ZnF(1-2-3)-BRCT_PARP1_ was ∼6 times lesser than PARP1 (Figure S8a & Figure 5a), which support previous structural studies showing that WGR_PARP1_ domain assists Zn-fingers for DNA binding^24^. On the other hand, binding affinities of ZnF(1-2-3)-BRCT_PARP1_ and PARP1 for SSB is almost same (Figure S8b and Figure 5b), which is in accordance with the reported data showing that only ZnFs, not WGR, are involved in SSB recognition^25^.

**Figure 5.**
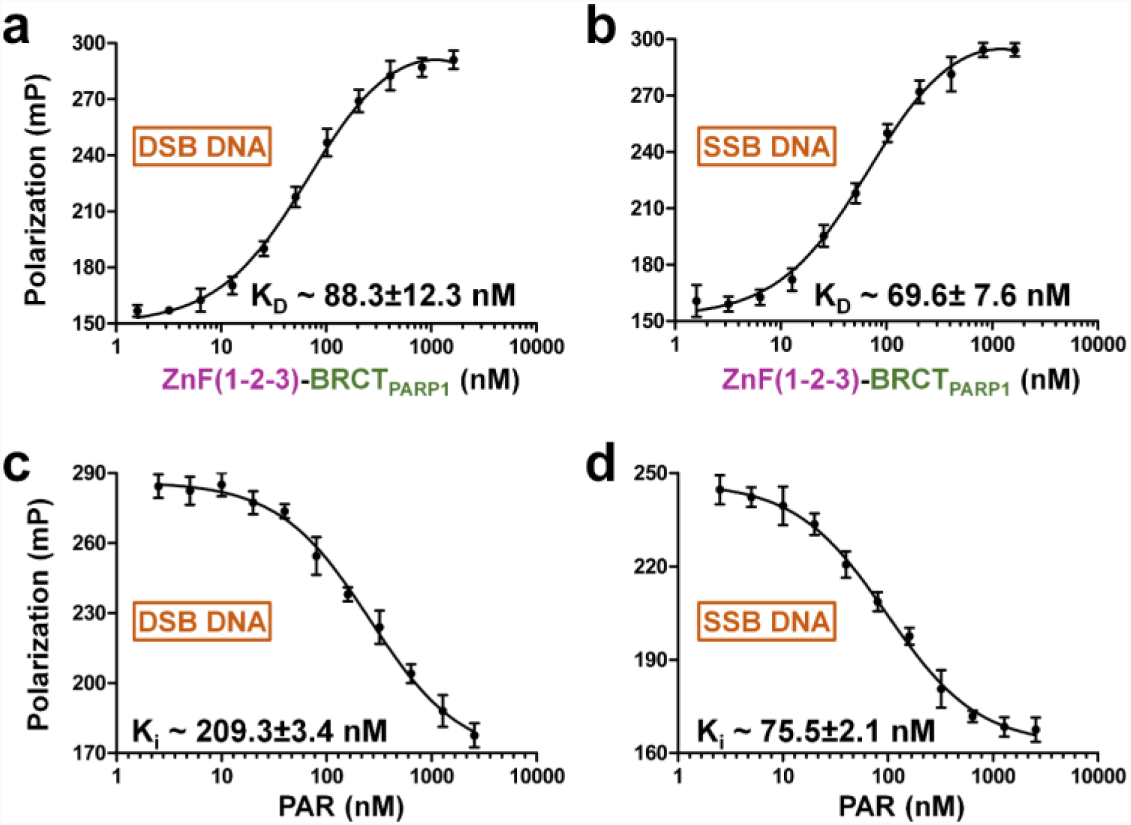
Fluorescence polarization binding and competitive binding studies of ZnF(1-2-3)-BRCT_PARP1_ to different DNA-breaks. FP binding studies of ZnF(1-2-3)-BRCT_PARP1_ with **a** DSB DNA and **b** SSB DNA. FP was calculated for increasing the concentration of ZnF(1-2-3)-BRCT_PARP1_ incubated with a fixed concentration of 5-FAM labeled DNA. K_D_ is reported as mean ± s.e.m. for three independent experiments. Competition binding studies of unlabeled PAR (increasing concentration) binding to ZnF(1-2-3)-BRCT_PARP1_ complexed with fluorophore labelled **c** DSB DNA and **d** SSB DNA. Competition binding studies were performed as described in Figure 4. The error bars indicate the standard deviation. All the data shown are from n=3 independent experiments. K_i_ values is reported as mean ± s.e.m.

Further, we performed FP based competition binding studies of PAR and DNA with the same variant of PARP1 (ZnF(1-2-3)-BRCT_PARP1_). The *K*_*i*_ of the unlabeled competitor PAR was determined to be ∼209 and ∼76 nM, respectively for DSB and SSB DNAs bound to ZnF(1-2-3)-BRCT_PARP1_ (Figure 5c-d). Comparative analyses show that (A) *K*_*i*_ of PAR for DNA displacement increased by a factor of ∼2 and ∼3, respectively, for DSB and SSB bound to ZnF(1-2-3)-BRCT_PARP1_ variant, compared to PARP1 (Figure 4, 5c-d). (B) Though PARP1 binds the above DNAs tighter than ZnF(1-2-3)-BRCT_PARP1_ (Figs. 5 a-b and Figure S8), lesser PAR concentration is required to dissociate DNA from PARP1 than the WGR lacking variant ZnF(1-2-3)-BRCT_PARP1_ (Figs. 4 and 5 c-d). Based on these results we conclude that the WGR domain might allosterically enhance the destabilization of PARP1-DNA complexes by binding to PAR.

These studies suggest that PAR triggers the release of DNA from PARP1 by directly competing with DNA to bind to WGR and ZnF3 domains, and possibly, induce global conformational changes in the PARP1 making it incompetent to recognize DNA by Zn-finger domains leading to dissociation of PARP1-DNA complex (Figs. 4 and 5 c-d). Adaptation of compact structural conformation from the extended beads-on-string conformation by PARP1 upon DNA binding^24^, supports our hypothesis of PAR-induced alternative global conformation in PARP1, though yet to be studied. It is noteworthy that certain PARP1 inhibitors (PARPi) bind to the catalytic domain of PARP1 and allosterically enhance either release or retention of PARP1 on the DNA^3,53^. PAR may function analogously to the PARPi(s), which release the PARP1 from the bound DNA, which needs further experimentation.

### PAR dependent catalytic stimulation of PARP1

Having established that PARP1 bind to PAR^46^ (Figure 1c), we sought to characterize the possibility of PAR modulating PARP1’s catalytic activity by performing SDS-PAGE based *in vitro* automodification assay and compared with DNA dependent PARylation. The disappearing PARP1 band and the appearance of smear above it (which shows an increase in the molecular weight due to PARylation) showed the PAR dependent stimulation of PARP1 catalytic activity (Figure 6a). We further quantified the extent of PARylation using PNC1-OPT assay which estimates the amount of nicotinamide released during the PARylation reaction. The amount of nicotinamide produced is equal to the NAD^+^ consumption by PARP1 and or ADPr units added. The assay uses the nicotinamidase (PNC1) enzyme to convert PARylation by-product nicotinamide to nicotinic acid with the release of ammonia. Further ammonia reacts with ortho-pthalaldehyde (OPT) to give fluorescence which is proportional to the catalytic activity of the PARP1^54^. The PNC1-OPT based *in vitro* PARylation enzyme activity assay shows a ∼20-fold increase in PARylation activity of PARP1 in the presence of PAR, compared to its absence after 30 minutes of incubation (Figure 6b). Therefore, like DNA breaks, PAR also stimulates the catalytic activity of PARP1 (Figure 6).

**Figure 6.**
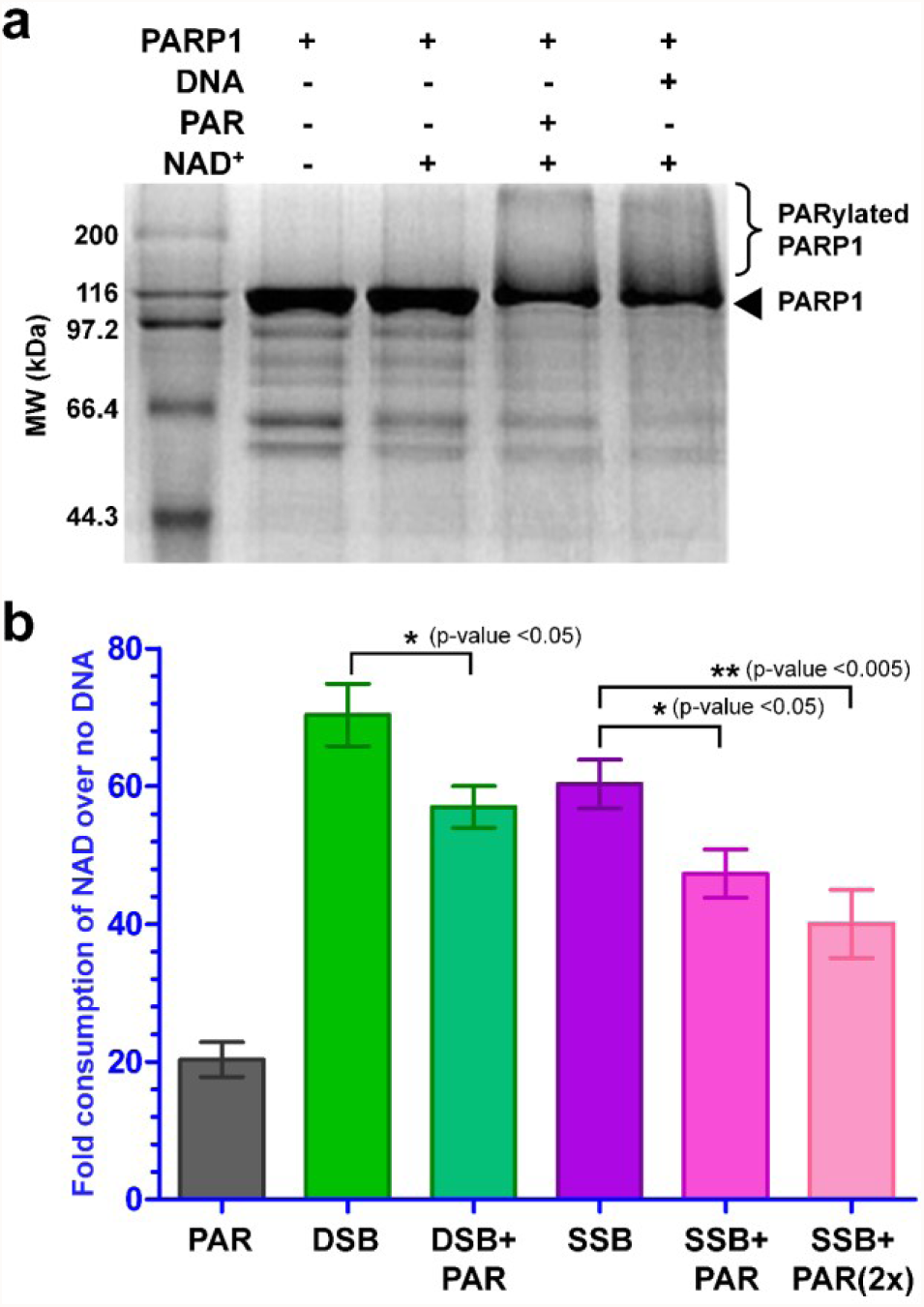
**a** SDS-PAGE automodification assay of PARP1 in presence of PAR and DNA. **b** Fluorometric assay of PARP1 PARylation activity in presence of PAR, different DNAs and DNA along with PAR plotted as fold consumption of NAD^+^ over the basal PARP1 activity^33^, i.e. in absence of DNA/PAR. PARylation activity of PARP1 was determined using fluorescence-based PNC1-OPT assay and fluorescence reading was measured at 30 minutes^61^. The measured fluorescence of the experimental set (with NAD^+^) was subtracted from the control (without NAD^+^) and the resultant is used to calculate the amount of NAD^+^ consumed (which is equal to ADP-ribose formed or the extent of PARylation). The p-values were calculated using an unpaired t-test. Error bars indicate the standard deviation of the n=3 independent experiments.

Since, our results on stimulation of the catalytic activity of PARP1 by PAR contradict the previously published data^39^, we have repeated these experiments with different batches of purified PARP1 and PAR. Each time our results of both PNC1-OPT and SDS-PAGE auto-modification assays are consistent. PARP2 catalytic activity stimulation by PAR supports our work^39^. This property of the PARP1 suggests that DNA independent, PAR-dependent, stimulation of enzymatic activity may be required for PARylation of proteins involved in nuclear processes other than DNA repair^37,55,56^. In this context, it is interesting to note that non-DNA dependent; RNA, protein, PAR (current study), etc. bind to different and/or overlapping regions of the PARP1, and stimulation of PARP1 catalytic activity is far less and/ or different from the DNA dependent stimulation^14,37,57^ (Figure S9), suggesting that these molecules fine tune the PARP1 activities for processes other than DNA damage signaling.

### Effect of PAR on DNA dependent stimulation of PARP1’s catalytic activity

Stimulation of PARP1 upon DNA-breaks recognition by the DNA binding domains is the hallmark of allosteric regulation of PARP1’s catalytic activity. Our studies show that different domains of PARP1 recognize PAR and DNA with differential binding affinity (Figure 2, 3 and S4). Here, we set out to investigate whether PAR binding to PARP1 can regulate the DNA dependent stimulation of PARP1, as it is already reported that DSB and SSB stimulate the PARP1 activity^24,25,33,35^. Our PNC1-OPT based activity assay shows that DSB and SSB DNA independently stimulate PARP1 activity by ∼70 and ∼60 folds, respectively (Figure 6b), compared to their absence after 30 minutes of incubation. Though PAR stimulates the catalytic activity of the PARP1, the PAR dependent stimulation is significantly less (3-3.5 times) compared to DNA dependent activity of the PARP1 (Figure 6b). Multiple residues on different regions of the PARP1 undergo PARylation upon DNA dependent stimulation, possibly all the residues may not be accessible for PAR dependent stimulation that might be a reason for less PAR dependent stimulation of PARP1 activity. At the same time, PAR decelerates the SSB and DSB DNA dependent stimulation of the PARP1’s catalytic activity by 18-24% (Figure 6b). Doubling the PAR concentration resulted in further reduction in SSB DNA stimulated PARP1’s catalytic activity suggesting lesser PARP1-DNA complexes with an increase in PAR concentration (Figure 6b). The results herein suggest that DNA dependent allosteric stimulation of PARP1’s catalytic activity is negatively regulated by PAR binding to PARP1.

Independent stimulation of the PARP1 catalytic activity by PAR and DNA (Figure 6), suggests that PAR and DNA could synergistically stimulate the PARP1 activity. Surprisingly, instead of enhancing, PAR decelerates the DNA dependent enzymatic activity of PARP1 (Fig. 6b). Possible explanation for this phenomenon is, given that PARP1 has overlapping DNA and PAR binding domains, and at a given point of time either PAR or DNA can bind to PARP1 and stimulates its catalytic activity. For PAR being an agonist, a stimulator of PARylation activity, the magnitude of retardation of DNA dependent PARP1 PARylation by PAR might be much more than what we observed in our experiments. In nutshell, PAR acts as both a partial allosteric stimulator and partial allosteric inhibitor (partial antagonist) of PARP1 (Figure 7). Recently, it has been reported that protein factor, HPF1, stimulates the DNA-dependent and independent self-PARylation of PARP1 and PARP2 as well as the trans-PARylation of histones in the complex with nucleosome^14^. Likewise, PAR stimulates the self-PARylation of PARP1 but, unlike HPF1, it hampers the DNA-dependent self-PARylation of PARP1. However, the effect of PAR or PARylated proteins on the trans-PARylation activity of PARP1 on histones, other proteins, or nucleic acid substrates of PARP1 needs to be addressed. Like Histone H4, PAR also stimulates PARP1’s catalytic activity through binding to its C-terminal domains^57^. However, it’s not known whether H4 hampers DNA dependent stimulation of PARP1’s activity like PAR.

**Figure 7.**
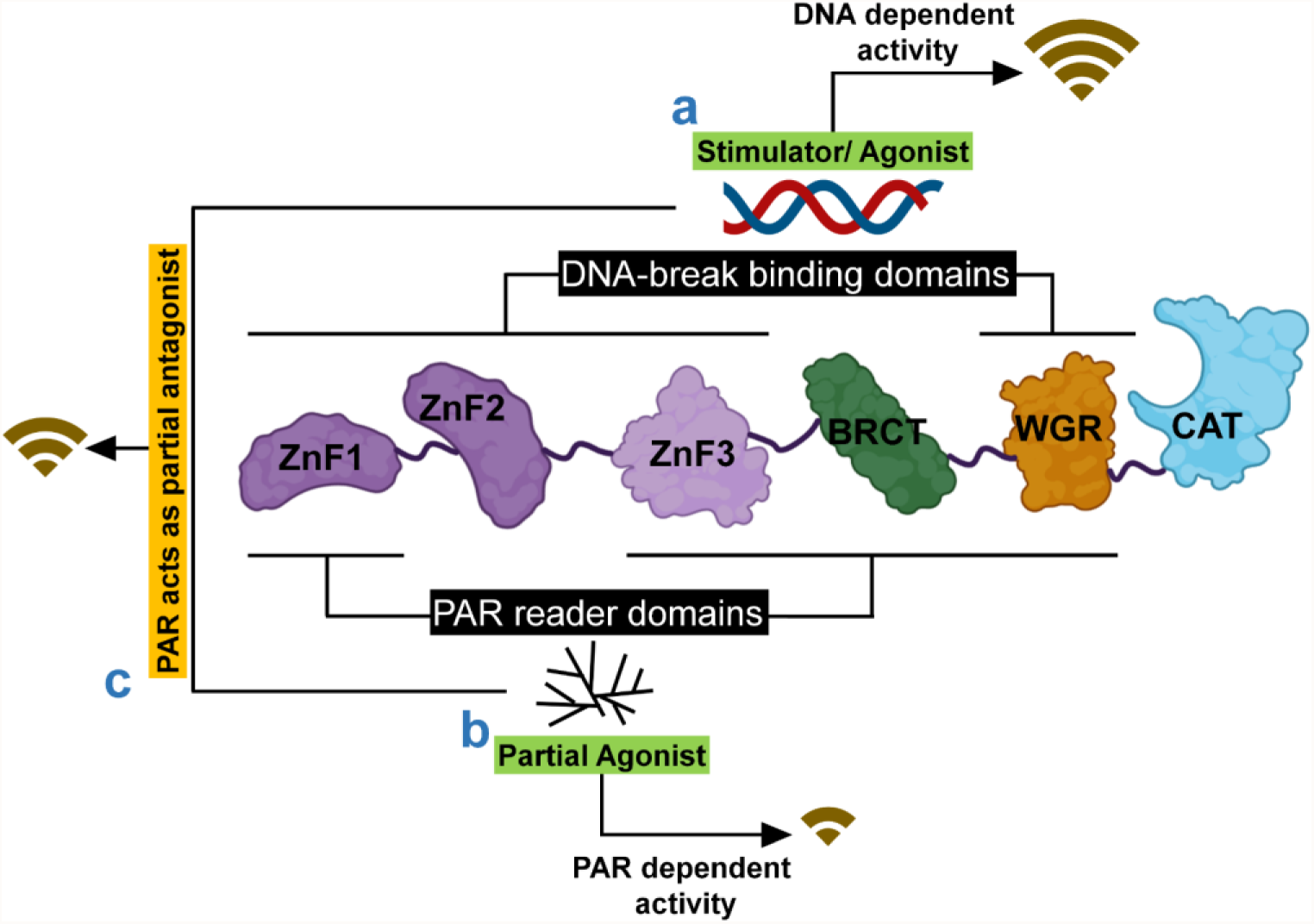
Schematic representation of proposed regulation of PARP1 activities by PAR and DNA breaks. **a** Recognition of DNA-break by DNA-break binding domains of PARP1 induces global conformational changes in the protein that leads to its maximal activation, thus DNA acts as a PARP1 stimulator/ agonist. **b** Binding of PAR, to PAR reader domains of PARP1 may lead to spatial reorganization of PARP1 domains to get into a certain conformation that stimulates the catalytic activity of PARP1, independent of DNA. **c** There are distinct yet overlapping domains for PAR and DNA breaks binding: Binding affinity of PARP1 domains to DNA breaks and PAR are ZnF1≈ZnF2>>ZnF3 and ZnF3≈WGR≈BRCT>>ZnF1>>ZnF2, respectively. Therefore, displacement of DNA by PAR from the DNA-PARP1 complex retards the DNA-dependent stimulation but at the same time PAR binding itself stimulates the PARP1 activity. Thus, the activation is modulated by cross-talk between DNA and PAR binding to PARP1.

## Conclusion

Our study revealed interesting features of PARP1: (i) PARP1 has distinct high-affinity PAR and DNA binding regions, in addition to overlapping low-affinity regions, (ii) PAR reader domains of PARP1 also recognize PARylated protein, (iii) PAR competes with DNA to bind to PARP1 and induce DNA dissociation, (iv) WGR domain plays an important role in PAR induced DNA dissociation from PARP1. We have also shown that PAR is a partial agonist as it stimulates PARP1 catalytic activity, though lesser than DNA dependent stimulation. It suggests an alternative pathway of stimulation of PARP1’s catalytic activity in the nucleus of a cell independent of DNA damage signaling. Our work unearths that PAR being a partial antagonist, decelerates the DNA-dependent stimulation of PARP1 activity. Further, structural studies of PAR with PAR binding domains of PARP1 (individual or in combination) and PARP1 would reveal mechanistic insights on the PAR dependent regulation of PARP1 and help in designing allosteric inhibitors or stimulators of PARP1. Moreover, these inhibitors would be very specific, since most of the available inhibitors (FDA approved or in clinical trials) target the NAD^+^ binding site, which could lead to non-specific targeting/ inhibition of other proteins that use NAD^+^ as a cofactor.

Given that PARP1 is subjected to allosteric regulation by the variety of molecules including activators and inhibitors^37,58,59^, further studies related to the cross-talk among allosteric ligands, that modulate the catalytic and DNA binding activities of PARP1 would reveal its yet-known roles and regulations in cellular systems. In addition, this work can be an example for unravelling the effect exerted by the combination of allosteric ligands (inhibitors and activators), bind to overlapping and/or distinct sites, on the variety of receptors or modular enzymes.

## Materials and Methods

### Cloning, expression and purification of PARP1, variants of PARP1, PARP2, PNC1 and OAS1

The full length human PARP1 cDNA cloned in pET28-a(+) was purchased from GenScript, USA. PARP1 (aa: 1-1014), ZnF(1-2-3)-BRCT_PARP1_ (aa: 1-494), and individual domain constructs were generated by cloning in bacterial expression vector pRSFDuet-1 vector between SacI and XhoI restriction sites. Domain length of individual constructs is same as given in Langelier et al., 2011^60^. *E. coli* Rosetta 2(DE3) were transformed with the constructs for expression. An hour before inducing the culture with 0.2 mM IPTG, 100 μM ZnSO_4_ was added. Culture was induced at 16°C for 16-18 h, then harvested. The pellet was resuspended in a lysis buffer containing 500mM NaCl, 50mM HEPES pH 7.5, 20mM Imidazole and 3mM β-mercaptoethanol. Protease inhibitor cocktail tablets (Pierce™ Protease Inhibitor, Thermo Scientific) were added in case of PARP1 only. The cell suspension was sonicated, the lysate was clarified by centrifugation and the proteins were purified in 3-steps. First, supernatant was passed through Ni-NTA affinity column (5ml, HisTrap column, GE Healthcare), followed by ion exchange Heparin column (HiTrap Heparin HP, GE Healthcare) with salt gradient of (250-1000mM NaCl for PARP1 and 50mM-1000mM NaCl for all other constructs) and finally through gel filtration (16/600 200pg or 16/600 75pg, GE Healthcare, depending on the size of the protein) column pre-equilibrated with buffer 150mM NaCl, 10mM HEPES, 2% Glycerol and 2mM β-mercaptoethanol). Purified proteins were concentrated, flash frozen and stored at -80°C.

The full length human PARP2 isoform 2 cloned in pET28-a(+) was generously gifted by Dr. John Pascal. PARP2 was expressed in *E. coli* BL21(DE3) at conditions same as PARP1 and purified using Ni-NTA affinity and ion-exchange chromatography using the protocol mentioned for PARP1.

PNC1 from Saccharomyces cerevisiae cloned in pET41-b(+) was obtained as a generous gift from Prof. Antonio Barrientos. *E. coli* Rosetta 2(DE3) was transformed with the construct for expressing the protein. PNC1 was expressed and purified as reported by Edwards et al. (2021)^61^.

Full-length human 2’-5’-oligoadenylate synthase 1 (OAS1) has been purchased from Genscript and cloned in the pRSF-Duet1 vector. Protein was expressed and purified using the same protocol as for PNC1.

### PAR generation, purification and fractionation

PAR was generated by performing automodification reaction of PARP1, cleaved from the protein and purified according to the protocol given in Lin et al. (2018)^62^. Obtained PAR was diluted with 20mM Tris-HCl buffer pH 7.5 and was loaded into Resource Q anion exchange 1ml column (GE Healthcare) for fractionation with a linear gradient of 0 - 1M NaCl at 0.25ml/min flow rate, with FPLC UV filter set at 254nm. Obtained separate fractions were precipitated by adding 1/10^th^ volume of 3M sodium acetate (pH 5.2), and 2.5 times volume of ice-cold 100% ethanol. After incubating overnight in -20°C, the precipitates were separated by centrifugation at 15000g for 30 min at 4°C. The pellet was washed with 1ml of 70% ethanol twice by centrifugation at 15000g for 10min. Pellet was air dried and resuspended in 20μl of autoclaved distilled water. 45 mer ssDNA would correspond to ∼25 mer PAR, since molecular weight of ADP ribose is ∼1.8 times of a nucleotide and charge on 2 nucleotides equals charge on one unit of ADP-ribose. Purified PAR fraction used for the binding studies is given in the Figure S3. Silver staining was performed to visualize the fractionated PAR on native PAGE. Concentration was calculated by taking the absorbance at 258nm and using the molar extinction coefficient of ADPr (13.5mM^-1^ cm^-1^).

### Generation of Biotinylated PAR

Biotin labeling of PAR was carried out as reported by Ando et al. (2019) with a few modifications^63^. 10 mM PAR was incubated with 40 mM biotin-14-dATP (Jena Biosciences), 50 mg/mL LMW poly(I:C) (Invivogen), 50 mg/mL OAS1 in 1x labeling buffer (20 mM TrisHCl pH 7.5, 20 mM Magnesium Acetate, 2.5 mM DTT) at 37°C for 2 h. Further poly(I:C) was removed purified PAR by RNase R treatment at 37°C for 1 h in presence of 100 mM KCl. Protein was removed by treating with 50 μg ml^−1^ proteinase K and 0.15% SDS for 30 min at 50°C followed by incubation for 3 min at 95°C. The samples were desalted on a Sephadex G-25 column (Amersham Biosciences) equilibrated with buffer containing 50mM NaCl and 10mM HEPES.

### Nucleic acids generation and annealing

HPLC purified lyophilized oligonucleotides and complementary strands (listed in Table S1) were purchased from oligo synthesis facility of W.M. Keck Foundation, Yale School of Medicine, USA and dissolved in a buffer containing 10 mM Tris-Cl (pH 7.5), 50 mM NaCl and 3 mM MgCl_2_. DSB DNA as prepared by mixing the complementary strands in an equimolar ratio. A dumbbell-forming 45-mer DNA oligonucleotide was used to make SSB DNA. SSB and DSB DNAs were annealed as previously reported^64^.

### Generation of PARylated PARP2

PARylated PARP2 was generated using the same automodification protocol as used for PARP1 with an incubation time of 30min, after which the reaction mixture was passed through the salt exchange column (HiPrep 16/10 Desalting, GE Healthcare) with FPLC UV filter set at 254nm and then concentrated using 10kDa concentrator. Concentration of PARylated PARP2 was calculated by subtracting the absorbance measured before and after PARylation reaction at 258nm and 280nm, using the molar extinction coefficient of ADPr.

### Bio-Layer Interferometry (BLI)

BLI-based binding kinetics studies were carried out on the OctetK2 instrument (Pall Fortebio). Biotinylated PAR was captured on High precision Streptavidin (SAX) Dip and Read™ biosensors (Pall Fortebio) with a threshold of 0.4-0.5 nm. For association kinetics measurement, the probes were dipped into wells containing varying concentrations of PARP1 (3-fold serial dilutions 500–12.2 nM for PAR) in buffer (150mM NaCl, 25mM Tris, 2% Glycerol and 3mM β-mercaptoethanol) for 120-150 s, followed by dipping into wells containing buffer for 200 seconds to measure the dissociation kinetics. For reference, one of the probes was dipped into buffer during the association and dissociation phases. The measurement was run at 25°C. The data were analysed and fit to a 1:1 model using ForteBio Data Analysis 12 (Pall Fortebio).

### Isothermal Titration Calorimetry (ITC) binding studies

ITC binding studies were carried out at 25°C using LV Affinity ITC (TA Instruments). The reference cell was filled with deionized water. For all binding studies unless stated otherwise, PAR (10-30 μM) or PARylated PARP2 (10-25 μM) were taken in cell (350μl) and PARP1 domains (0.2-1 mM) were taken in syringe. For ZnF1_PARP1_-PAR and ZnF2_PARP1_-PAR titrations 250 µM PAR was used in cell. 100 µM PAR was taken in cell for CAT_PARP1_-PAR titration. 20 consecutive injections of 2.5 μl at an interval of 120s were performed while stirring the cell content at 125 rpm. As control, buffer was titrated against PAR (15 µM). For sample preparation, titrants and titrands were diluted in gel filtration buffer (150mM NaCl, 10mM HEPES, 2% Glycerol and 2mM β-mercaptoethanol). All the binding studies were repeated 3 times. Analysis and processing of the binding isotherm was performed using NanoAnalyse software provided by TA instruments, and the molar dissociation constant (K_D_) values are reported as mean±sem (n=3).

### Fluorescence Polarization (FP) studies

For fluorescence polarization DNA binding studies, samples were prepared in a buffer containing 150 mM NaCl, 50mM HEPS, 2% Glycerol and 2mM β-mercaptoethanol. Fluorophore, 5-FAM, labelled DNAs (Table S1) were used for binding and competition binding studies. For DNA binding studies with PARP1 and ZnF(1-2-3)-BRCT_PARP1_, 20 nM fluorophore labelled DNA was used and protein concentration was varied from 1.6 nM to 3.2 µM. For binding studies of individual zinc fingers, 10 nM of labelled DNA and, 12 nM to 12.5 µM (ZnF1_PARP1_/ZnF2_PARP1_/ZnF3_PARP1_) were used. For the competition binding assay, 40 nM of protein was incubated with 20 nM of probe DNA and unlabeled PAR was added in varying concentrations of 1.5 nM to 5 µM (PARP1 and ZnF(1-2-3)-BRCT_PARP1_) at room temperature for 30 min. The fluorescence polarization data were collected on SpectraMax ID5 (Molecular devices). The FP values were plotted against the variable entity into a nonlinear regression model. The *K*_*i*_ were calculated from Binding-Competitive model in GraphPad Prism v5. The reported *K*_*i*_ and K_D_ represent a mean±s.e.m. of three independent experiments.

### SDS-PAGE based auto-modification assay of PARP1

To perform PARP1 auto-modification assay, PARP1 (11.3 µM) in auto-modification buffer was incubated with DNA (1 µM) or PAR (5 µM) for 10 min^60^. Following the incubation, 0.2 mM NAD^+^ was added to the reaction. The auto-modification reaction was kept at room temperature for 60 min and quenched by the addition of SDS-loading dye. The sample was analyzed on 12% SDS-PAGE stained with coomassie brilliant blue.

### PNC1-OPT assay

PNC1-OPT assay was performed to measure the amount of NAD^+^ consumed during PARylation by PARP1, as described by Edwards et al (2021)^61^. To measure activation by PAR, 1µM PARP1, 5 µM PAR and 0.2 mM NAD^+^ were used. To determine the effect of PAR on DNA-dependent PARylation activity of PARP1, the reaction was performed with using 1 µM PARP1, 2µM DNA and/or 5/10 µM PAR and 0.2 mM NAD^+^ The fluorescence was measured using the plate reader (EnSpire™ Multimode Plate Reader by PerkinElmer, Inc.) with monochromators set to excitation at 420 nm and emission at 450 nm and the fluorescence data were analyzed for PARylation activity as reported previously^54^. Then fold of NAD+ consumption in the presence of PAR, DNA or PAR & DNA over absence of PAR and DNA was calculated^33^ The above data analysis was performed from three independent reactions. The p-values were calculated using an unpaired t-test. Error bars indicate standard deviation of three independent experiments.

No unexpected or unusually high safety hazards were encountered.

## Acknowledgments

This work was supported by the ‘Research and Development’ scheme under ‘Basic Research in Modern Biology’ of Department of Biotechnology (DBT), Government of India. E.R. thanks the DBT and the Science and Engineering Research Board, Department of Science and Technology, Government of India for the Ramalingaswami re-entry fellowship and Early Career Research Award, respectively. W.D. thanks University Grants Commission, Government of India for the fellowship. S.A. thanks the Ministry of Human Resource Development, Government of India for the fellowship. Authors thank Dr. Jyotsnendu Giri, Biomedical Engineering Department, IIT Hyderabad for providing access to multimode reader. Authors also thank Dr. John Pascal, Thomas Jefferson University, Philadelphia, USA and Prof. Antonio Barrientos, University of Miami, FL, USA for providing the pET28-a(+)-PARP2iso2 plasmid and pET41-b(+)-PNC1 plasmid, respectively.

## Author Information

### Contribution

E.R. conceived, designed and supervised the research. W.D. and S.A. designed and performed the research. W.D., S.A., and E.R. analysed the data. W.D. and E.R. wrote the manuscript.

