## Supplementary Information for "PAR recognition by PARP1 regulates DNA-dependent activities and independently stimulates catalytic activity of PARP1"

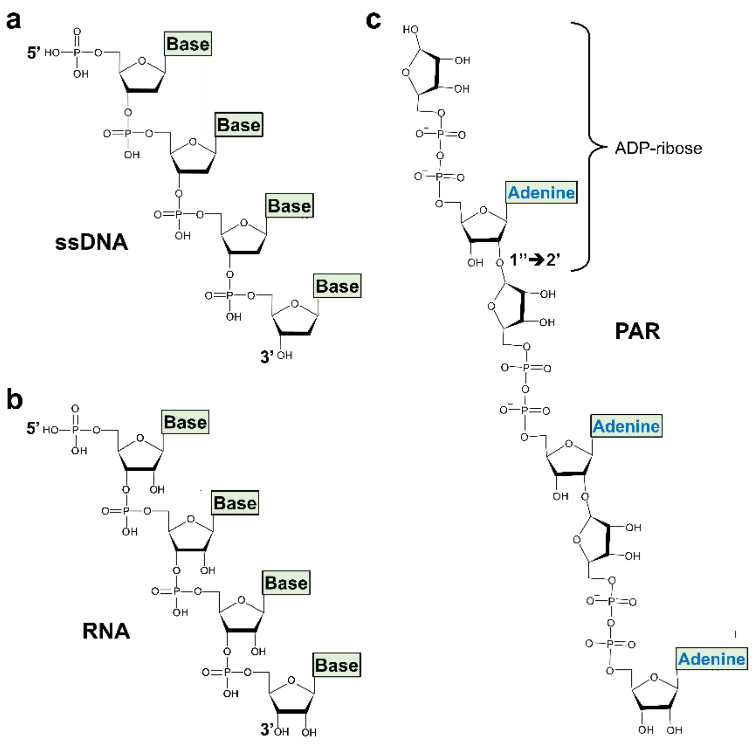


**Figure S1. 2D Schematic representations. a** DNA where base could be A/T/G/C, **b** RNA where base could be A/U/G/C. **c** PAR which has repeating units of ADP-ribose.

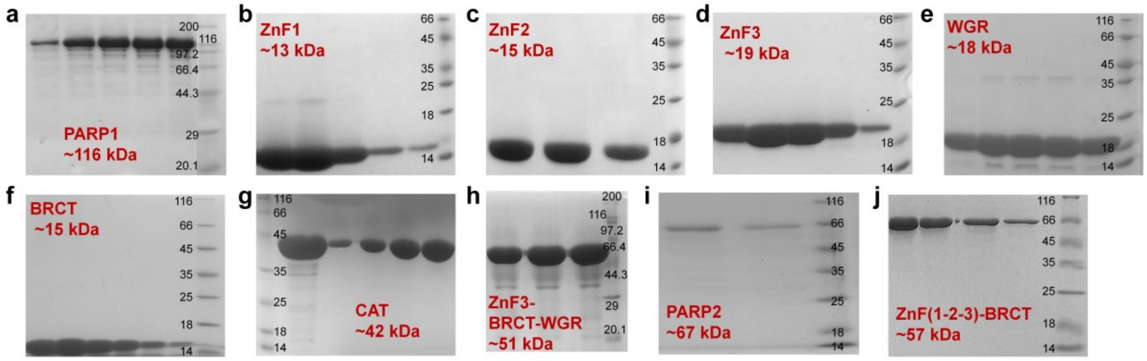


**Figure S2. SDS-PAGE of gel filtration purified individual or combination of domains of PARP1 used in the study.**

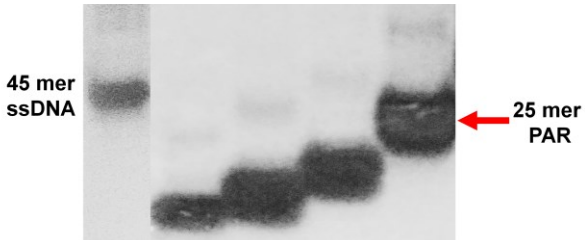


**Fig S3. Native-PAGE of fractionated PAR obtained after anion exchange chromatography.** The leftmost lane shows the 25 mer ssDNA band, other lanes show PAR of different lengths.

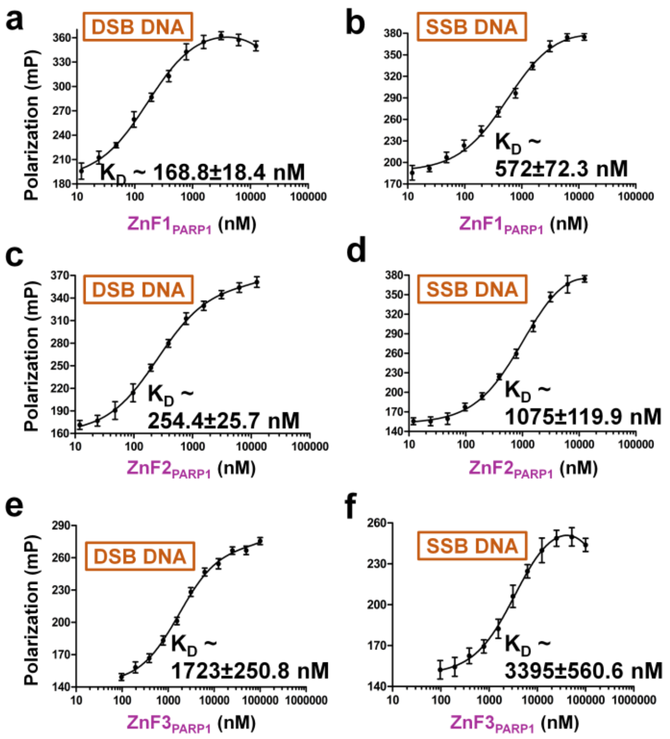


**Figure S4.** FP binding studies of DSB and SSB DNAs binding to **(a-b)** ZnF1$_{PARP1}$, **(c-d)** ZnF2$_{PARP1}$, and **(e-f)** ZnF3$_{PARP1}$. FP was calculated for increasing the concentration of protein incubated with the fixed concentration of 5-FAM labeled DNA. The error bars indicate the standard deviation. $K_D$ is reported as mean±s.e.m. of three independent experiments.

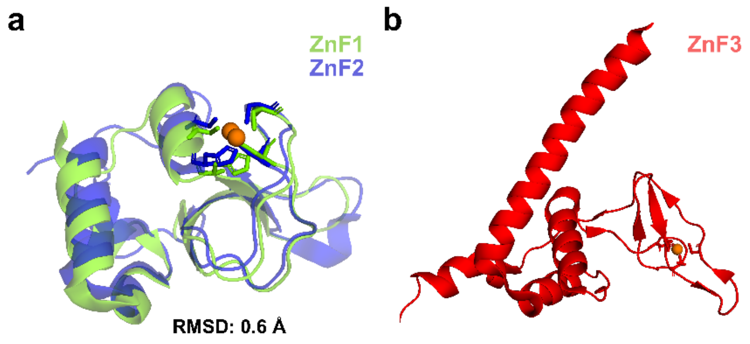


**Figure S5. Structural comparison between ZnFs of PARP1. a** Structural superposition of $ZnF1_{PARP1}$ (PDB ID: 3OD8) on $ZnF2_{PARP1}$ (PDB ID: 3ODC). **b** Structure of $ZnF3_{PARP1}$ (PDB ID: 2RIQ) is different from $Zn1_{PARP1}$ and $ZnF2_{PARP1}$, therefore could not be superposed on $ZnF1_{PARP1}$ or $ZnF2_{PARP1}$.

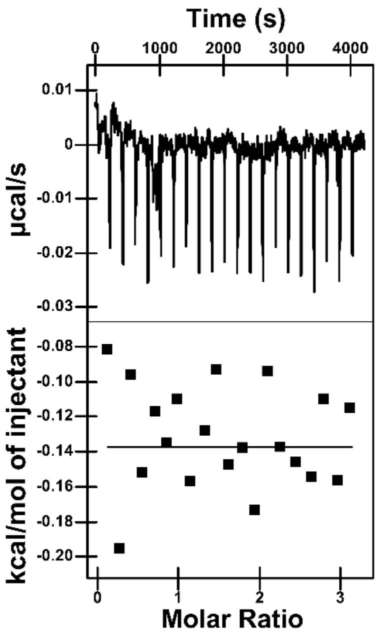


**Figure S6. ITC thermogram of Buffer-PAR titration.**

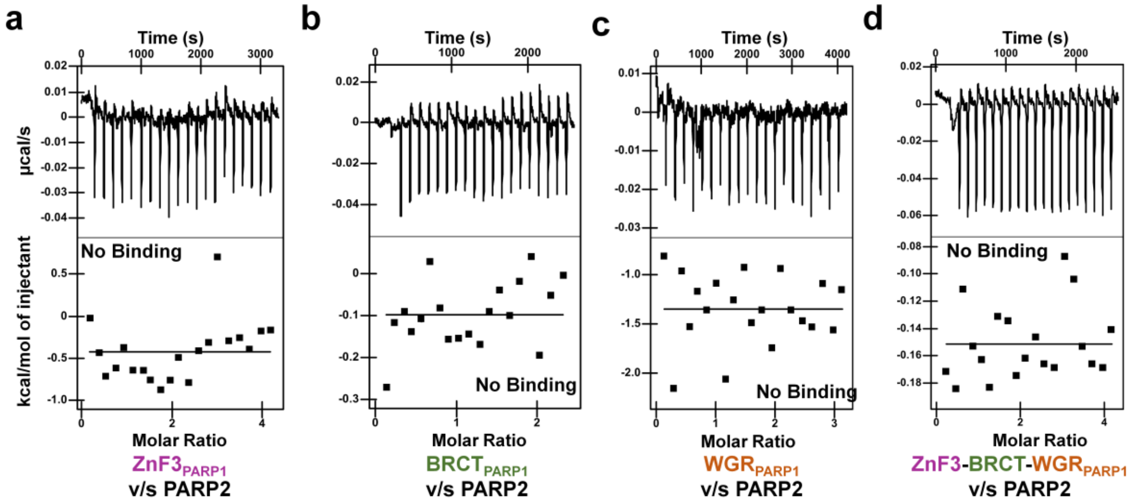


**Figure S7. Binding studies of PARP2 with domains of PARP1.** ITC measurement of PARP2 titrated with **a** $ZnF3_{PARP1}$, **b** $BRCT_{PARP1}$, **c** $WGR_{PARP1}$ and **d** ZnF3-BRCT-$WGR_{PARP1}$.

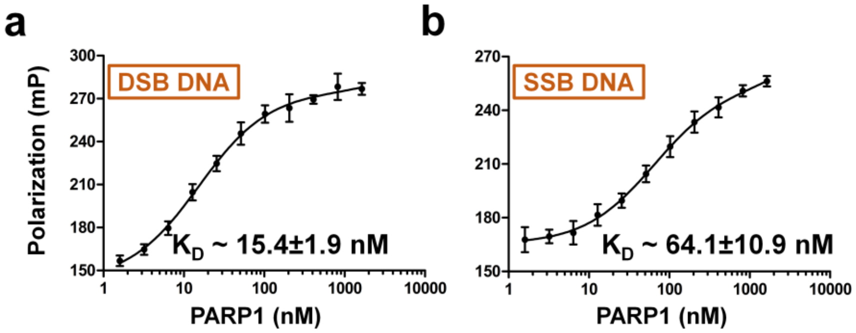


**Figure S8. FP-based DNA binding studies of PARP1**. Binding of PARP1 to **a** DSB DNA, and **b** SSB DNA. The error bars indicate the standard deviation. $K_D$ is reported as mean±s.e.m. of three independent experiments.

### Supplementary Table 1. List of DNA used in the study.

| DNA type | Sequence |
|---|---|
| **DSB DNA** | 5’-GCCTACCGGTTCGAATGGCAGC-3’<br>3’-CGGATGGCCAAGCTTACCGTCG-5’ |
| **SSB DNA** | 5’-P-<br>GCTGGCTTCGTAAGAAGCCAGCTCGCGGTCAGCTTGCTGACCGCG-3’<br>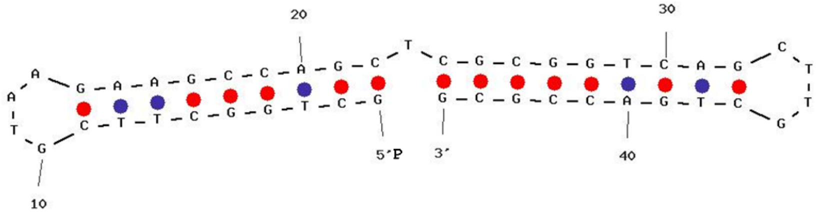 |
| **5-FAM labelled DNA** | |
| **DSB DNA** | 5’-GCCTACCGGTTCGAATGGCAGC-3’<br>3’-CGGATGGCCAAGCXTACCGTCG-5’ |
| **SSB DNA** | 5’-P-<br>GCTGGCTXCGTAAGAAGCCAGCTCGCGGTCAGCTTGCTGACCGCG -3’<br>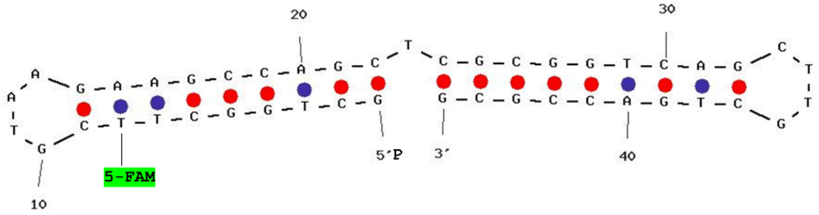 |

P= Phosphate; X = 5-FAM-dT
